# Fast and flexible design of novel proteins using graph neural networks

**DOI:** 10.1101/868935

**Authors:** Alexey Strokach, David Becerra, Carles Corbi-Verge, Albert Perez-Riba, Philip M. Kim

## Abstract

Protein structure and function is determined by the arrangement of the linear sequence of amino acids in 3D space. Despite substantial advances, precisely designing sequences that fold into a predetermined shape (the “protein design” problem) remains difficult. We show that a deep graph neural network, ProteinSolver, can solve protein design by phrasing it as a constraint satisfaction problem (CSP). To sidestep the considerable issue of optimizing the network architecture, we first develop a network that is accurately able to solve the related and straightforward problem of Sudoku puzzles. Recognizing that each protein design CSP has many solutions, we train this network on millions of real protein sequences corresponding to thousands of protein structures. We show that our method rapidly designs novel protein sequences and perform a variety of *in silico* and *in vitro* validations suggesting that our designed proteins adopt the predetermined structures.

**One Sentence Summary:** A neural network optimized using Sudoku puzzles designs protein sequences that adopt predetermined structures.

## Introduction

Protein structure and function emerges from the specific geometric arrangement of their linear sequence of amino acids, commonly referred to as a fold. Engineering novel protein sequences has a broad variety of uses, including academic research, industrial process engineering (*1*), and most notably, protein-based therapeutics, which are now a very important class of drugs (*2, 3*).

However, despite extraordinary advances, designing a sequence from scratch to adopt a desired structure, referred to as the “inverse folding” or “protein design” problem, remains a challenging task. Conventionally, a sampling technique such as Markov-chain Monte Carlo is used to generate sequences optimized with respect to a force-field or statistical potential (*3*–*5*). Limitations of those methods include the relatively low accuracy of existing force fields (*6, 7*) and the inability to sample more than a miniscule portion of the vast search space (sequence space size is 20^N^, N being the number of residues). While there have been successful approaches that screen many thousands of individual designs using *in vitro* selection techniques (*8, 9*), those approaches remain reliant on labor-intensive experiments.

Here we overcome those limitations by combining a classic idea with a novel methodology. Filling a specific target structure with a new sequence can be formulated as a constraint satisfaction problem (CSP) where the goal is to assign amino acid labels to residues in a polymer chain such that the forces between interacting amino acids are favorable and compatible with the fold. To overcome previous difficulties with phrasing protein design as a CSP (*10, 11*), we elucidate the rules governing constraints using deep learning. Such methods have been applied to a vast diversity of fields with impressive results (*12*–*14*), partly because they can infer hitherto hidden patterns from sufficiently large training sets. For proteins, the set of different protein folds is only modestly large with a few thousand superfamilies in CATH (*15*). Indeed, previous attempts at using deep learning approaches for protein design used structural features and thus only trained on relatively small datasets and achieved moderate success as of yet without any experimental validation (*16*–*20*). However, the number of sequences that share these structural templates is many orders of magnitude larger (about 70,000,000 sequences map to the CATH superfamilies), reflecting the fact that the protein design problem is inherently underdetermined with a relatively large solution space. Thus, a suitable deep neural network trained on these sequence-structure relationships could potentially outperform previous models to solve the protein design problem.

The distance matrix is commonly used to represent protein folds (*21*). It is an NxN matrix consisting of the distances between residues, optionally restricted to only interacting pairs of residues that are within a certain distance of one another. The distance matrix can be thought of as placing constraints on pairs of residues, such that the forces governing the interaction between those residues are not violated (e.g. interactions between residues with the same charge or divergent hydropathicities are usually not well tolerated). A given protein structure, corresponding to a single distance matrix, can be formed by many different homologous sequences, and those sequences all satisfy the constraints imposed by the distance matrix. Such solutions to this constraint satisfaction problem (CSP) are given to us by evolution and are available in sequence repositories such as Pfam (*22*) or Gene3D (*23*). While the rules of this CSP for a specific protein fold can be found by comparing sequences from one of such repositories and can be captured as Hidden Markov Models (HMM) or position weight matrices (PWMs), often represented as sequence logos, it has thus far not been possible to deduce general rules—those that would connect any given protein fold or distance matrix with a set of sequences. Here, we use a graph neural network, denoted ProteinSolver, to elucidate those rules. The graph in this case is made up of nodes, corresponding to amino acids, and edges between those nodes, corresponding to the spatial interactions between amino acids, as represented in the distance matrix. The edges thus represent the constraints that are imposed on the node properties (amino acid types).

We show that a ProteinSolver network trained to elucidate the rules governing the CSP of protein folding by reconstructing masked sequences shows remarkable success in generating novel protein sequences for a predetermined fold. Previous approaches to protein design were hampered by the enormous computational complexity, in particular when taking into account backbone flexibility. Our approach sidesteps this problem and delivers plausible designs for a wide range of folds. We expect that it would also be able to generate sequences for completely novel imagined protein folds. Furthermore, as a neural network approach, its evaluation is many orders of magnitude faster than classical approaches and should enable the exploration of vastly more potential backbones.

Finally, we present a web server which allows users to run a trained ProteinSolver model to generate sequences matching the geometries of their own reference proteins. We hope that this web server will lower the barrier to entry for protein design and will facilitate the generation of many novel proteins. The web server is freely available at: http://design.proteinsolver.org.

## Results and Discussion

### Network architecture

As there had been little previous work in using neural networks to solve CSPs (*24, 25*), we first had to devise a network architecture that would be well-suited for this problem. In order to facilitate this search, we focused on designing a neural network capable of solving Sudoku puzzles, which is a well-defined CSP (*25*) for which predictions made by the network can easily be verified. We treat Sudoku puzzles as graphs having 81 nodes, corresponding to squares on the Sudoku grid, and 1701 edges, corresponding to pairs of nodes that cannot be assigned the same number (Fig. 1B). The node attributes correspond to the numbers entered into the squares, with an additional attribute to indicate that no number has been entered, the edge indices correspond to the 1701 pairs of nodes that are constrained such that they cannot have the same number, and the edge attributes are all the same value because all edges in the graph impose identical constraints. We generated 30 million solved Sudoku puzzles using the sugen program (*26*), which first generates a solved Sudoku grid using a backtracking grid filler algorithm, and then randomly removes numbers from that grid until it generates a Sudoku puzzle with a unique solution at the requested difficulty level. Neural networks with different architectures were trained to reconstruct the missing numbers in the Sudoku grid, by minimizing the cross-entropy loss between predicted and actual values. Throughout training, we tracked the accuracy that those networks achieve on the training dataset (Fig. 2A, blue line) and on the validation dataset (Fig. 2A, orange line), which contains 1000 puzzles which were excluded from the training dataset.

**Fig. 1.**
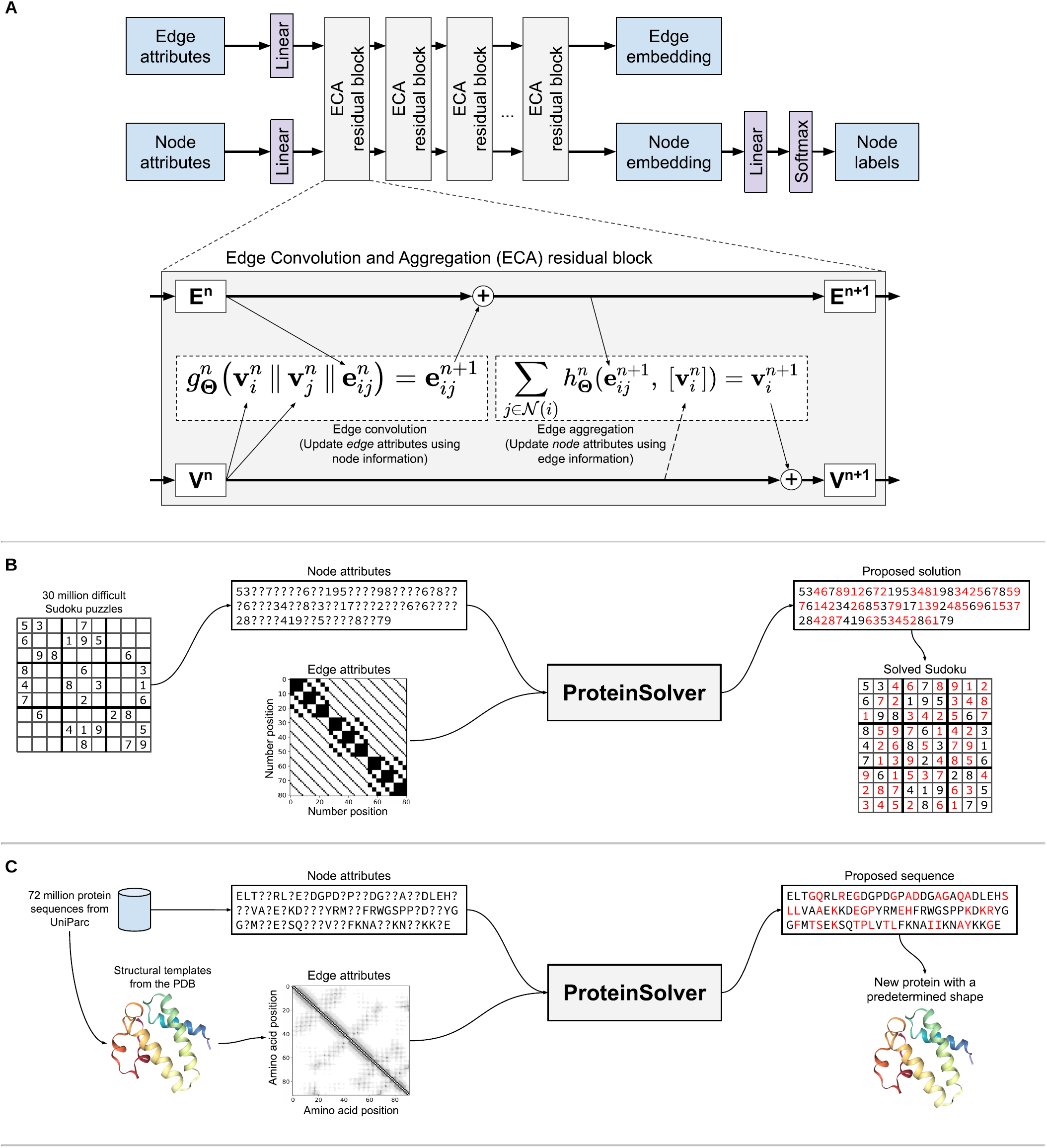
Graph convolutional neural network used by ProteinSolver to assign node labels that satisfy the provided node and edge constraints. **(A)** ProteinSolver network architecture. **(B)** Training a ProteinSolver network to solve Sudoku puzzles. Node attributes encode the numbers provided in the starting Sudoku grid. Edge attributes encode the presence of constraints between pairs of nodes (i.e. that a given pair of nodes cannot be assigned the same number). **(C)** Training a ProteinSolver network to reconstruct protein sequences. Node attributes encode the identities of individual amino acids. Edge attributes encode Euclidean distances between amino acids and the relative positions of those amino acids along the amino acid chain.

**Fig. 2.**
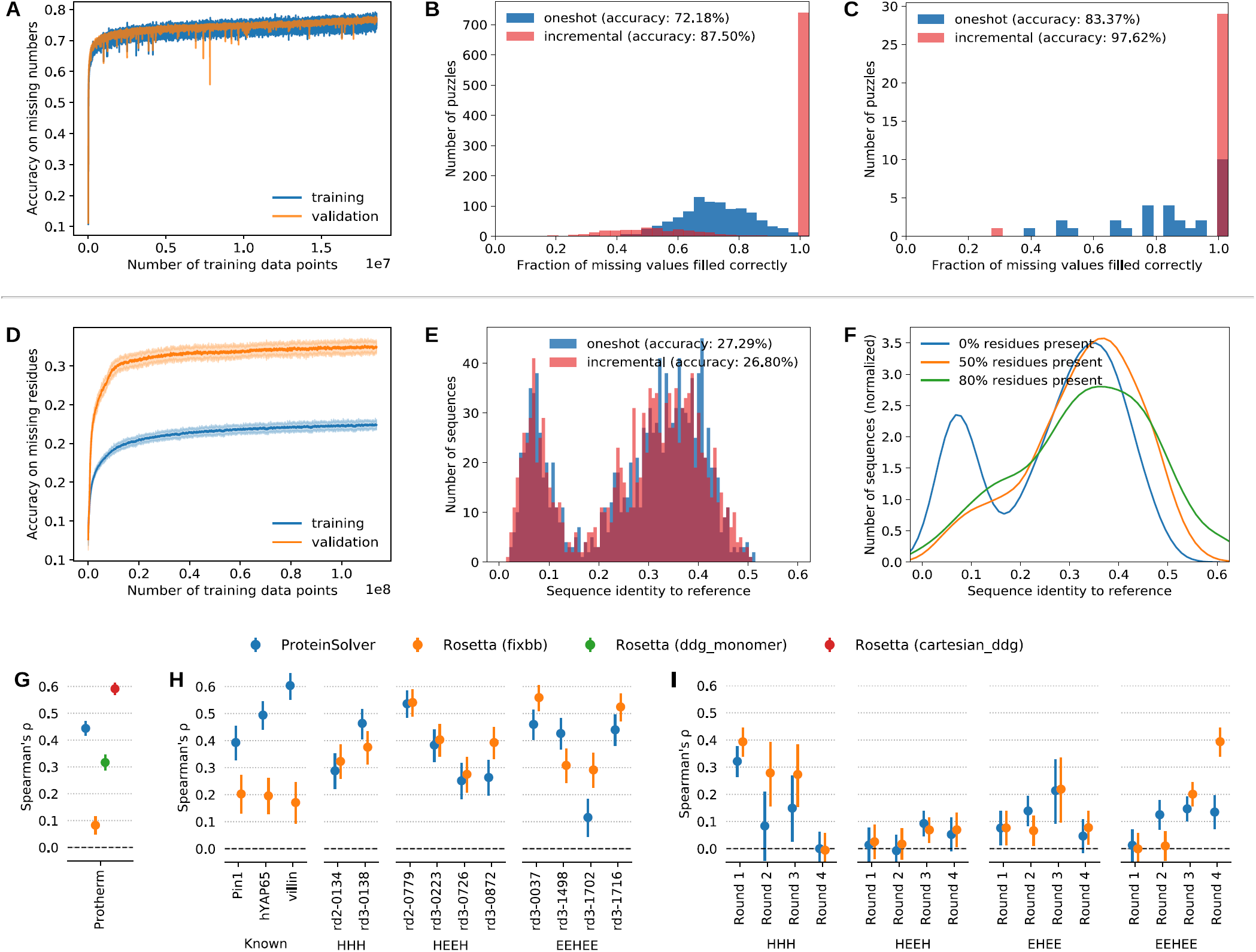
**(A)** Training and validation accuracy of the ProteinSolver network being trained to solve Sudoku puzzles. **(B-C)** Accuracy achieved by the ProteinSolver network, trained to solve Sudoku puzzles, on the validation dataset **(B)**, comprised of 1000 Sudoku puzzles generated in the same way as the training dataset, and on the test dataset **(C)**, comprised of 30 Sudoku puzzles extracted from an online Sudoku puzzle provider. Predictions were made using either a single pass through the network (blue bars) or by running the network repeatedly, each time taking a single prediction in which the network is the most confident (red bars). **(D)** Training and validation accuracy of the ProteinSolver network being trained to recover the identity of masked amino acid residues. During training, 50% of the amino acid residues in each input sequence were randomly masked as missing. **(E)** Accuracy achieved by the ProteinSolver network on the test dataset, comprised of 10,000 sequences and adjacency matrices of proteins possessing a different shape (Gene3d domain) than proteins in the training and validation datasets. In the case of blue bars, predictions were made using a single pass through the network, while in the case of red bars, predictions were made by running the network repeatedly, each time taking a single prediction in which the network is the most confident. **(F)** Sequence identity between generated and reference sequences in cases where 0%, 50%, or 80% of the reference sequences are made available to the network. **(G)** Spearman correlation coefficients between experimentally-measured changes in protein stability associated with mutation and predictions made using Proteinsolver (blue), Rosetta’s fixbb protocol (orange), Rosetta’s ddg_monomer protocol (green), and Rosetta’s cartesian_ddg protocol (red). **(H)** Spearman correlation coefficients between changes in protein stability associated with mutation, measured using an enzyme digestion assay, and predictions made using Proteinsolver (blue) and Rosetta’s fixbb protocol (orange). **(I)** Spearman correlation coefficient between the stability of proteins designed *de novo* using Rosetta to match a specific architecture, and predictions made using Proteinsolver (blue) and Rosetta’s fixbb protocol (orange). HHH, HEEH, EHEE, and EEHEE denote proteins designed *de novo* using Rosetta in order to have a helix-helix-helix, helix-sheet-sheet-helix, sheet-helix-sheet-sheet, or sheet-sheet-helix-sheet-sheet architecture, respectively.

After a broad scan over different neural network architectures that we conceived for this problem, we converged on the “ProteinSolver” graph neural network architecture presented in Fig. 1A. The inputs to the network are a set of node attributes and a set of edge attributes describing interactions between pairs of nodes. The node and edge attributes are embedded in an *m*-dimensional space using linear transformations or a multi-layer perceptron. The resulting node and edge attribute embeddings are passed through N residual edge convolution and aggregation (ETA) blocks. In the convolution step, we update the edge attributes using a modified version of the edge convolution layer (*27*), which takes as input a concatenation of node and edge attributes and returns an update to the edge attributes. In our work, *g*_Θ_ is a multi-layer perceptron, although other neural network architectures are possible. In the aggregation step, we update node attributes using an aggregation over transformed edge attributes incident on every node. In our work, *h*_Θ_ is a learned linear transformation, although other neural network architectures, including attention layers (*28*), are possible.

In the case of Sudoku, the fully-trained network with optimized hyperparameters predicts correctly 72% of the missing numbers in a single pass through the network and close to 90% of the missing number if we pass the input through the network multiple times, each time adding as a known value a single prediction from the previous iteration in which the network is the most confident (Fig. 2B). Similar accuracy is achieved on an independent test set containing puzzles from an online Sudoku puzzle provider (Fig. 2C).

### Reconstructing and evaluating protein sequences

After optimizing the general network architecture for the well-defined problem of solving Sudoku puzzles, we applied a similar network to protein design, which is a less well-defined problem than Sudoku and for which the accuracy of predictions is more difficult to ascertain (Fig. 1C). We treat proteins as graphs, where nodes correspond to the individual amino acids and edges correspond to shortest distances between pairs of amino acids, considering only those pairs of amino acids that are within 12 Å of one another. The node attributes specify the amino acid, with an additional flag to indicate that the amino acid is not known, while the edge attributes include the shortest distance between each pair of amino acids in Cartesian space and the number of residues separating the pair of amino acids along the protein chain.

We compiled a dataset of 72 million unique Gene3D domain sequences from UniParc (*29*) for which a structural template could be found in the PDB (*30*), and we trained the network by providing as input a partially-masked amino acid sequence together with the adjacency matrix adapted from the structural template, and minimizing the cross-entropy loss between network predictions and the identities of the masked amino acid residues (Fig. 1C). The training and validation accuracies achieved by the network reach a plateau after around 100 million training examples, with a training accuracy of ∼22% and a validation accuracy of ∼32%, when half of the residues are masked in the starting sequence (Fig. 2D-E). The training accuracy is lower than the validation accuracy because, while the training dataset has no restriction on the similarity between sequences and structural templates, for the validation dataset we included only those sequences for which a structural template with at least 80% sequence identity to the query could be found. Reconstruction accuracy is considerably lower than Sudoku (Fig. 2A-C), as was expected: the Sudoku CSP has a single well-defined solution, thus an accuracy approaching 1 is possible. By contrast, each protein adjacency matrix can be adopted by many different sequences, and the achieved accuracy of 30%-40% roughly corresponds to the common level of sequence identity within a protein fold (*15*). Evaluating our network by having it reconstruct sequences from our validation dataset, we observe a bimodal distribution in sequence identities between generated and reference sequences, with a smaller peak around 7% sequence identity and a larger peak around 37% sequence identity, corresponding to generated sequences about as similar to the reference sequences as other members of their family (see Fig. 2F). Note that reconstruction accuracy substantially higher than what we achieve would likely be an artifact, as it is already common for sequences with the achieved level of sequence identity to adopt the same fold and thus fulfill the same CSP.

We next asked whether the score assigned by a trained network to single mutations can be predictive of whether those mutations are stabilizing or destabilizing. We speculated that a destabilizing mutation would also disrupt some of the constraints in the CSP and would thus be scored unfavourably by the graph neural network. We find that predictions made using ProteinSolver, which is trained solely to reconstruct protein sequences, show a significant correlation with experimentally measured changes in protein stability reported in Protherm (*31*) (Spearman ρ: 0.44; p < 0.001) and, in fact, show a stronger correlation than predictions made using Rosetta’s fixbb and ddg_monomer protocols (Fig. 2G). While predictions made using ProteinSolver show a weaker correlation than predictions made using Rosetta’s cartesian_ddg protocol, both the cartesian_ddg protocol, and the beta_nov16 energy function used by that protocol, have been optimized in sight of the Protherm dataset and may have inadvertently overfit on this data (*32*). Furthermore, Rosetta’s cartesian_ddg protocol performs extensive structural relaxation and sampling around the site of the mutation and takes on the order of minutes to hours to evaluate a single mutation, while ProteinSolver can typically evaluate a mutation in under a second.

While the ProteinSolver network was not trained using any mutation data, we did not explicitly exclude the proteins in the Protherm dataset from our training dataset. In order to ascertain that the correlation between predictions made using ProteinSolver and the experimentally-measured ΔΔG values is not biased by the presence of the wild-type sequences in our training dataset, we calculated the correlation between predictions made using ProteinSolver and the effect of mutations on the stability of a number of proteins designed *de novo* to have a contrived shape (*9*) (Fig. 2H). While none of those *de novo* designs appear in the ProteinSolver training dataset, ProteinSolver nevertheless achieves similar correlations as Rosetta’s fixbb protocol for mutations in those proteins. Note that we did not make predictions for this dataset using Rosetta’s ddg_monomer and cartesian_ddg protocols because of the heavy computational resources that would be required and the fact that evaluating the effect of multi-residue mutations is not explicitly supported by those protocols.

Finally, in order to evaluate how well ProteinSolver can score entire protein sequences and prioritize them for subsequent experimental evaluation, we calculated the correlation between the scores assigned by ProteinSolver to complete novel proteins that have been designed *de novo* using Rosetta, and the stability of those proteins, as measured using a high-throughput sequencing approach (*9*) (Fig. 2I). We observe that, for most protein geometries and rounds of selection, the correlation between scores assigned by ProteinSolver and the stability of the proteins is similar to the correlation observed for scores produced by the Rosetta protocol used to generate the protein library. One exception is the round 4 library of the EEHEE designs, where Rosetta achieves a Spearman correlation of ∼0.4 while ProteinSolver achieves a Spearman correlation of ∼0.14, although it should be noted that ProteinSolver achieves a significantly higher correlation for the round 2 library of designs with the same geometry.

### Generating new protein sequences with predefined geometries

Motivated by the observation that the ProteinSolver network is able to reconstruct protein sequences with a reasonable level of accuracy, and to assign probabilities to individual residues, as well as entire proteins, that correlate well with experimental measurements of protein stability, we sought to use the network to generate entire novel protein sequences for specific protein folds. To that end, we chose four protein folds that had been left out of the training set and cover the breadth of the CATH hierarchy, and for each of those folds, we extracted a distance matrix from a protein structure representative of that fold. We designed new protein sequences matching those distance matrices by starting with an entirely empty or “masked” protein sequences and, to each of the positions in those sequences, iteratively assigning residues by sampling from the residue probability distributions defined by the network (see Methods). This corresponds to *de novo* protein design—designing novel sequences for a given fold.

In total, we generated over 600,000 sequences for each of the four selected folds: serum albumin (mainly α; Fig. 3), alanine racemase (α+β; Supp. Fig. S1), PDZ3 domain (mainly β; Supp. Fig. S2), and immunoglobulin (mainly β; Supp. Fig. S3). The generated sequences tend to fall within 37-44% sequence identity with the native sequence (Fig. 3B; Supp. Fig. S1B, S2B, S3B), in line with the sequence identity within the respective protein families. Furthermore, the sequence conservation profiles of the generated sequences show that the network has a clear preference for specific residues at some positions, and in cases where several residues are equally preferred, those residues tend to have similar chemical properties (Fig. 3D; Supp. Fig. S1D, S2D, S3D). As no information about the sequences was provided to the network, it suggests that the network is able to learn the features pertinent to mapping structural information to sequences from the training dataset.

**Fig. 3.**
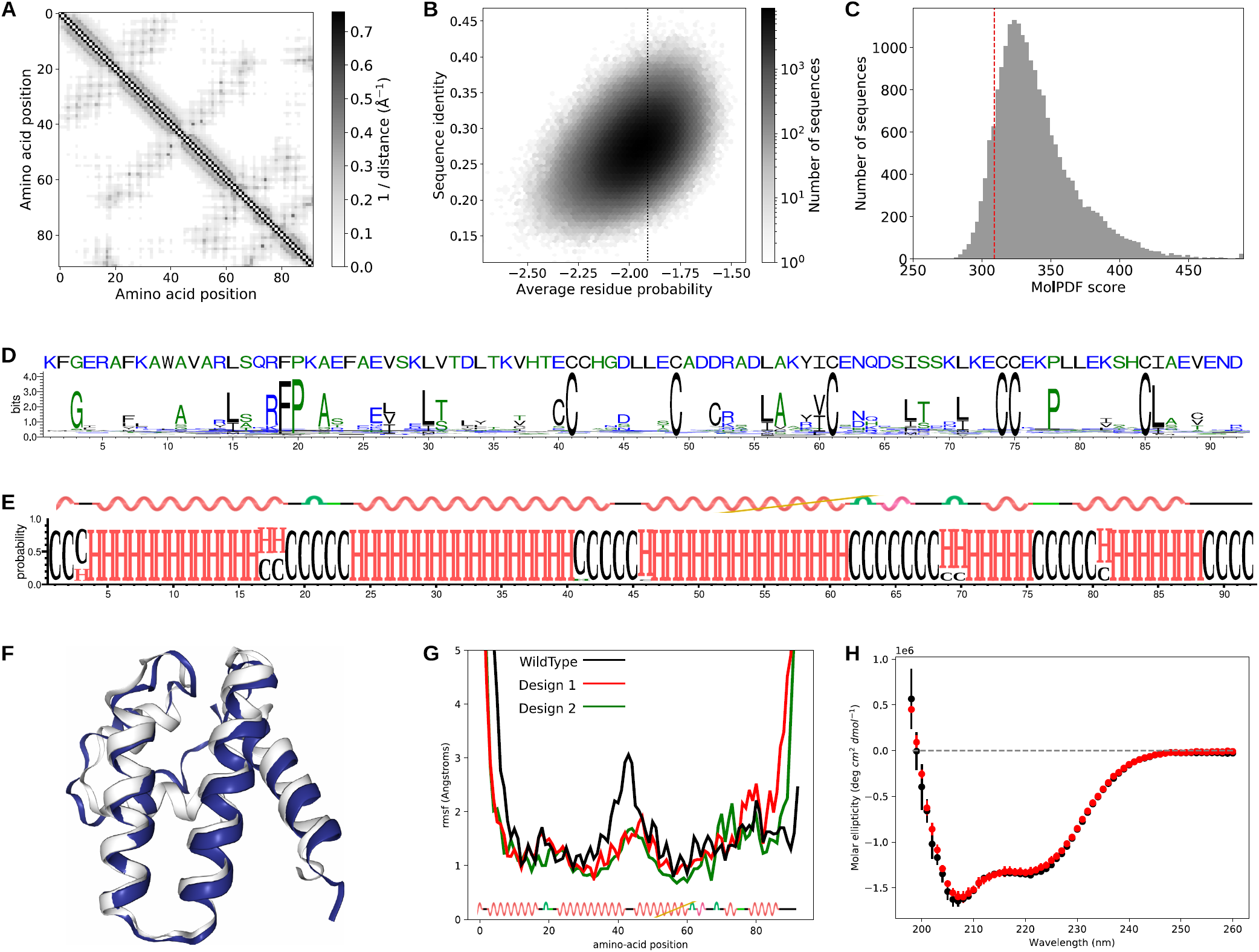
Computational and experimental validation of sequences generated to match the architecture of a *serum albumin*. **(A)** Amino acid contact map extracted from PDB structure *1N5U* and provided as input to the network. **(B)** Correlation between residue probabilities and sequence identities to the reference structure, for ∼2 million sequences generated by the trained ProteinSolver network. **(C)** Distribution of Modeller MolPDF scores for homology models constructed for a sample of 20,000 generated sequences. **(D)** Sequence LOGO showing the conservation of residues in the generated sequences. **(E)** Secondary structure LOGO showing predicted secondary structures for a sample of 20,000 generated sequences. **(F)** Structure of the reference protein (white) overlaid with the structure of a model produced for one of the generated sequences using QUARK, a de-novo structural modelling tool (blue). **(G)** Average residue fluctuation in 100 ns molecular dynamics simulations of the reference structure and two homology models of generated sequences. **(H)** Circular dichroism spectra of the reference protein (black) and a generated protein (red).

For each of the four selected folds, we selected the top 20,000 (10%) sequences, as scored by our network, and performed further computational validation. First, we used PSIPRED (*33*) to predict the secondary structure of each of the generated sequences (Fig. 3E; Supp. Fig. S1E, S2E, S3E), and we found that the predicted secondary structures of our designs match the secondary structures of the reference proteins almost exactly. Next, we created homology models of our designs, and we evaluated those homology models using a number of metrics, including the Modeller molpdf score (*34*) and the Rosetta REU score (*35*) (Fig. 3C; Supp. Fig. S1C, S2C, S3C). In all cases, the scores obtained for the designs are in the same range or better than the scores obtained for the reference structures, suggesting that the sequences that our network generated *do novo* indeed fold in the shape corresponding to the provided distance matrix. We also used QUARK (*36*), a *de novo* (not template-based) structure prediction algorithm to obtain structures for our sequences (Fig. 3F; Supp. Fig. S1F, S2F, S3F). For all four folds, the obtained structures match the reference structure almost exactly. Finally, we performed 100 ns molecular dynamics (MD) simulations of the reference structures and the homology models of our de novo designs (Fig. 3G; Supp. Fig. S1G, S2G, S3G). In all four cases, the designs show comparable fluctuation in molecular dynamics to the reference proteins, indicating that they are of comparable stability as the reference proteins and are thus stably folded.

After obtaining encouraging results from our computational validation experiments, for two of the four selected folds, we chose sequences with the highest combined network and Modeller NormDOPE score, and we attempted to express and evaluate those sequences experimentally. We had each of the sequences synthesized as oligonucleotides, and we expressed them as his-tagged constructs for Ni-NTA affinity purification. After purification, we evaluated the secondary structure of each protein using circular dichroism spectroscopy (CD). Each secondary structure element has a distinctive absorbance spectrum in the far-UV region, and thus similar folds should present similar absorbance spectra. For the two folds, serum albumin and alanine racemase, the selected sequences show a CD spectrum that is similar to the spectrum obtained for the native protein (Fig. 3E; Supp. Fig. S1E). This is particularly striking in the case of the serum albumin template, where the spectra are indistinguishable (Fig. 3E). The sequence generated for the alanine racemase template displayed a considerable loss of solubility compared to the sequence from the target structure. Although this made its characterization challenging, we were able to obtain clear spectra by combining a low ionic strength buffer (10 mM Na-phosphate, pH 8) with a 10 mm cuvette. While the resulting CD spectrum is somewhat different from the target (Supp. Fig. S1E), this may be due to technical issues resulting from low solubility or a more dynamic nature of the designed protein (consistent with the molecular dynamics in Supp. Fig. S1D). The spectrum definitely corresponds to a folded helical structure consistent with the predetermined fold. Taken together with the rest of the evidence from molecular dynamics and Modeller and Rosetta assessment scores, this strongly suggests that those generated sequences adopt the fold specified to the neural network.

## Conclusion

In this article, we present ProteinSolver, a graph neural network-based method for solving the protein design problem, formulated as a CSP, and generating protein sequences which satisfy the specified geometric and amino acid constraints. We show that a trained ProteinSolver network can reconstruct protein sequences with a high degree of accuracy, and that it assigns probabilities to individual residues, and to entire protein sequences, that correlate well with the stability of the resulting proteins. Finally, a trained ProteinSolver network can generate novel protein sequences which, according to extensive computational and experimental validation, fold into the same shapes as the reference proteins from which the geometric constraints are extracted.

There has been growing interest in using neural network-based approaches for protein representation learning and design (*19, 37*–*40*), with several new methods reported during the preparation of this manuscript (*41, 42*). While most methods are accompanied by a variety of metrics which attempt to illustrate the accuracy of the predictions, it is inherently difficult to evaluate the quality of generative models, as their ultimate goal is to generate entirely novel sequences with no existing counterparts. To address this concern, we have synthesised and experimentally validated several of our designs, and have shown that the designed proteins fold into stable structures with circular dichroism spectra that are consistent with their target shapes. As far as we are aware, ProteinSolver is currently the only machine learning based model whose predictions have undergone this level of experimental validation.

One limitation of existing methods for protein design is the steep learning curve and the high degree of domain expertise that is necessary to make reasonable predictions. We circumvent those limitations by developing a web server which the users can use to generate, in near-real time, hundreds to thousands of sequences matching a given protein topology and amino acid constraints (Supp. Fig. S4). It can be freely accessed at: http://design.proteinsolver.org. We believe that this web server will make our graph neural network-based approach to protein design accessible to the widest possible audience and will facilitate the generation of many novel proteins.

## Materials and Methods

### Data preparation

We downloaded from UniParc (*37*) a dataset of all protein sequences and corresponding domain definitions, and we extracted from this dataset a list of all unique Gene3D domains. We also processed the PDB database and extracted the amino acid sequence and the distance matrix of every chain in every structure. The distance matrix consists of distances between all pairs of residues that are within 12 Angstroms of one another, considering both the backbone and the sidechain residues in this calculation. Finally, we attempted to find a structural template for every Gene3D domain sequence, and we transferred the distance matrices from the structural templates to each of those sequences. The end result of this process is a dataset of 72,464,122 sequences and adjacency matrices, clustered into 1,373 different Gene3D superfamilies. We split this dataset into a training subset, containing sequences of 1,029 Gene3D superfamilies, a validation subset, containing sequences of 172 Gene3D superfamilies, and a test subset, containing sequences of another 172 Gene3D superfamilies. Instructions on how to download the training and validation datasets are provided on the ProteinSolver documentation page (https://ostrokach.gitlab.io/proteinsolver). A list of all resources used to construct those datasets is provided Supp. Table S1.

In the case of Sudoku, the training and validation datasets were generated using the sugen program (*26*) with the target difficulty of the puzzles set to 500. The training dataset was composed of 30 million generated puzzles, while the validation dataset was composed of 1000 puzzles that do not appear in the training dataset. The test dataset was comprised of 30 Sudoku puzzles collected from http://1sudoku.com (*43, 44*).

### Network implementation

The source code for ProteinSolver is freely available at https://gitlab.com/ostrokach/proteinsolver. The network was implemented in the Python programming language using PyTorch (*45*) and PyTorch Geometric (*46*) libraries. The repository also includes Jupyter notebooks that can be used to reproduce all the figures presented in this manuscript.

### Network architecture

We used Sudoku as a toy problem while optimizing the general design of the ProteinSolver network, including selecting the objective function that is optimized (masking a fraction of node labels and minimizing the cross-entropy loss between predicted and actual labels), tuning the specific implementation of edge convolutions and aggregations (using a 2-layer feed-forward network to update edge attributes, summing over linearly-transformed edge attributes of the incident edges to update node attributes, etc.), and selecting the types of non-linearities and normalizations that are applied (ReLU and LayerNorm, respectively).

Once we had a network that showed promising results in its ability to solve Sudoku puzzles, we tuned the specific hyperparameters of that network and selected variants that achieved the highest accuracies on the validation datasets. For the task of solving Sudoku puzzles (Fig. 2A-C), the model that achieved the highest accuracy on the validation dataset had 16 residual edge convolution and aggregation (ECA) blocks and a node and edge embedding space of 162. For the task of reconstructing protein sequences (Fig. 2D-F), the model that achieved the highest accuracy on the validation dataset had 4 residual blocks and a node and edge embedding space of 128.

### Scoring existing protein sequences

We evaluated the plausibility of existing protein sequences by using a trained ProteinSolver network to calculate the log-probability of every residue in those sequences, given all other residues, and taking the average of those log-probabilities. In effect, for every residue in a given sequence, we replaced the node label corresponding to that residue with a “mask” token, and we used a trained network to obtain the log-probability of the correct residue at the masked position. The score for the protein (e.g. Fig. 2I) was calculated as the average of the log-probabilities assigned to all residues. The score for a mutation (e.g. Fig. 2G,H) was calculated as the difference between log-probabilities of the mutant and the wild-type protein.

### Generating novel protein sequences

In order to calculate the most probable protein sequence, given a specific distance matrix, we evaluated two different approaches: one-shot generation and incremental generation. In one-shot generation, we passed the inputs with the missing node labels through the network only once, for every node accepting the label assigned by the network the highest probability. In incremental generation, we passed the inputs through the network once for every missing label. At each iteration, we accepted the label for which the network was making the most confident prediction, and we treated that label as given in all subsequent iterations. The one-shot generation method is substantially faster, requiring O(1), rather than O(N), passes through the network, while the incremental generation method appears to produce more accurate results, especially in the case of Sudoku (see Fig. 2C,E).

In order to generate *a library* of protein sequences, given a specific distance matrix, we used an approach similar to the incremental generation method described above. However, at each iteration, instead of deterministically accepting the residue to which the network assigns the highest probability, we select the residue by randomly sampling from the probability distribution assigned to that position by the network.

### Molecular dynamics

All water and ion atoms were removed from the structure with PDB codes 1N5U, 4Z8J, 4UNU, and 1OC7, corresponding to an all-α protein, a mainly-β protein, an all-β protein and a mix-αβ protein, respectively. The structural models for the generated sequences were produced using Modeller, with the PDB files described above serving as templates. Using TLEAP in AMBER16 (*47*) and AMBER ff14SB force field (*48*), the structures were solvated by adding a 12 nm^3^ box of explicit water molecules, TIP3P. Next, Na+ and Cl-counter-ions were added to neutralize the overall system net charge, and periodic boundary conditions were applied. Following this, the structures were minimized, equilibrated, heated over 800 ps to 300 K, and the positional restraints were gradually removed. Bonds to hydrogen were constrained using SHAKE (*49*) and a 2 fs time step was used. The particle mesh Ewald (*50*) algorithm was used to treat long-range interactions.

### Web server implementation

In order to make our method accessible to the widest possible audience, we developed a web server which allows the user to run a trained ProteinSolver model to generate new protein sequences matching geometric constraints extracted from a reference structure and, optionally, specific amino acid constraints imposed directly by the user (Supp. Fig. S4; http://design.proteinsolver.org). For the curious reader, we have also made available a web server which can be used to run a ProteinSolver model trained to solve Sudoku puzzles (http://sudoku.proteinsolver.org). The web servers were implemented using the Voila web framework and are deployed as Docker containers running on Google Kubernetes Engine. We use Knative in order to automatically scale the number of virtual machines and running Docker containers, providing a responsive experience for an arbitrary number of users while minimizing the overall cost of operation. The protein design web server is deployed to virtual machines equipped with an NVIDIA Tesla T4 GPU, with at most eight concurrent users sharing a single GPU.

### Experimental validation

#### Protein purification

All constructs were synthetized by Invitrogen GeneArt Synthesis (ThermoFisher) as Gene Strings. The construct designs included flanking BamHI and HindIII restriction sites for subcloning into a N-terminal 6xHis-tagged pRSet A vector. All genes were codon optimized for expression in *E. coli* using the GeneArt suite.

The pRSET A constructs were transformed into chemically competent E. coli OverExpress C41(DE3) cells (Lucigen) by heat shock and plated on LB-Carbenicillin plates. Colonies were grown in 15 ml of 2xYT media containing Carbenicillin (50 μg/mL) at 37 °C, 220 rpm until the optical density (O.D.) at 600 nm reached 0.6. Cultures were then induced with IPTG (0.5 mM) for 3h at 37°C. Cells were pelletized by centrifugation at 3000 g (4 °C, 10 min) and resuspended in 1 ml of BugBuster Master Mix (Millipore). The lysate mixtures were incubated for 20 min in a rotating shaker and the insoluble fraction was removed by centrifugation at 16000 g for 1 min. For a 15 ml preparation, 200 μl of Ni-NTA Agarose (QIAGEN) were pre-washed in Phosphate-buffered saline (PBS) and added to the supernatant from the cell lysate for 20 min at 4 °C in batch. The bound beads were washed thrice with PBS (1 mL) containing 30 mM of imidazole to prevent nonspecific interaction of lysate proteins. Proteins were eluted using PBS buffer with 500 mM imidazole and dialyzed against PBS (*51*).

Protein concentration was calculated using the Pierce BCA Protein Assay Kit (ThermoFisher). The concentration of 4beu_Design was corroborated by absorbance at 205 nm as described by Anthis et. al. (*52*). 1n5u and 1n5u_Design and 4beu eluted solubly at concentrations around 200-400 ug/ml. 4beu_Design presented a limit of solubility at around 15 ug/ml. Sample purity was assessed through Coomassie staining on SDS-PAGE with 4–20% Mini-PROTEAN TGX Precast Protein Gels, 10-well (Bio-Rad) and Mass Spectrometry (ESI).

#### Circular dichroism

All CD measurements were made on a Jasco J-810 CD spectrometer in a 1 mm Quartz Glass cuvette for high performance (QS) (Hellma Analytics) with the exception of 4beu_Design where a 10 mm Quartz Glass cuvette (QS) (Hellma Analytics) was preferred. 1n5u and 1n5u_Design were analysed in PBS whereas 4beu and 4beu_Design were analysed in 10 mM Na-Phosphate, pH 8. The CD spectra were collected in the 198 nm to 260 nm wavelength range using a 1 nm bandwidth and 1 nm intervals at 50 nm/min; each reading was repeated then times. All measurements were taken at 20 °C.

## Acknowledgments

We thank Quaid Morris for valuable discussions.

## Author contributions

Conceptualization: AS and PMK. Data curation: AS and DB. Formal analysis and Methodology: AS. Software: AS and DB. Resources: AS, DB, CC, APR and PMK. Validation: AS, CC, DB and APR. Writing: AS, DB, CC, APR and PMK. Project administration: PMK. Supervision: PMK. Funding acquisition: PMK.

## Data and materials availability

All code and data are publicly available at: https://gitlab.com/ostrokach/proteinsolver.

## Supplementary Materials (see separate file)

**Tables S1-S5**

**Figures S1-S5**

## Supplemental Material

### Supplemental Tables

**Table S1.**
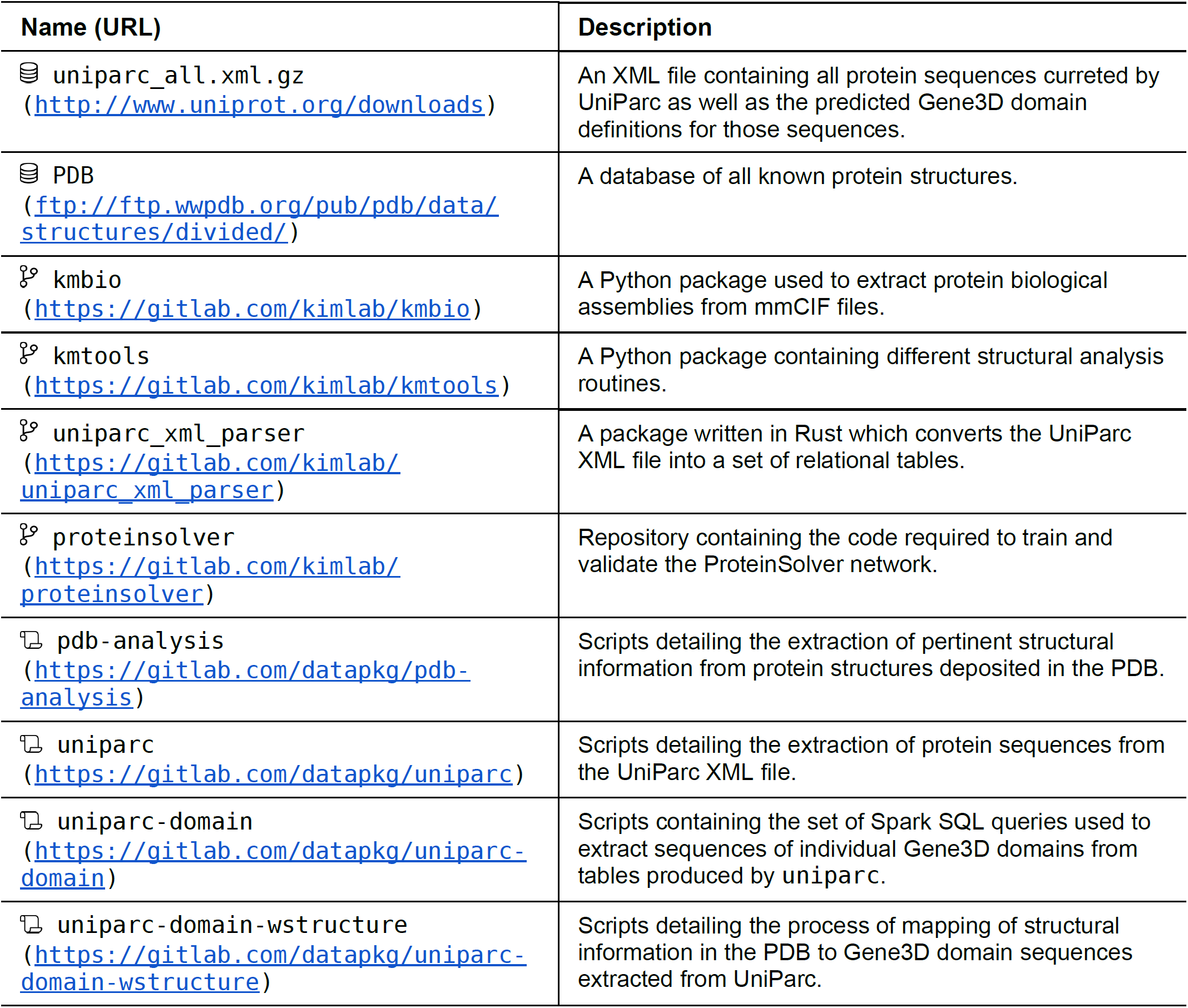
Resources used to prepare the training and validation datasets required to train the ProteinSolver network for protein design. Resources denoted with a database icon 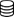 correspond to input data sources. Resources denoted with a git branch icon 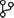 correspond to software packages used in the preparation of the datasets. Resources denoted with a scroll icon 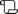 correspond to repositories containing Jupyter notebooks detailing the individual steps of data processing.

| Name (URL) | Description |
| --- | --- |
| uniparc_all.xml.gz<br>(http://www.uniprot.org/downloads) | An XML file containing all protein sequences curated by UniParc as well as the predicted Gene3D domain definitions for those sequences. |
| PDB<br>(ftp://ftp.wwpdb.org/pub/pdb/data/structures/divided/) | A database of all known protein structures. |
| kmbio<br>(https://gitlab.com/kimlab/kmbio) | A Python package used to extract protein biological assemblies from mmCIF files. |
| kmtools<br>(https://gitlab.com/kimlab/kmtools) | A Python package containing different structural analysis routines. |
| uniparc_xml_parser<br>(https://gitlab.com/kimlab/uniparc_xml_parser) | A package written in Rust which converts the UniParc XML file into a set of relational tables. |
| proteinsolver<br>(https://gitlab.com/kimlab/proteinsolver) | Repository containing the code required to train and validate the ProteinSolver network. |
| pdb-analysis<br>(https://gitlab.com/datapkg/pdb-analysis) | Scripts detailing the extraction of pertinent structural information from protein structures deposited in the PDB. |
| uniparc<br>(https://gitlab.com/datapkg/uniparc) | Scripts detailing the extraction of protein sequences from the UniParc XML file. |
| uniparc-domain<br>(https://gitlab.com/datapkg/uniparc-domain) | Scripts containing the set of Spark SQL queries used to extract sequences of individual Gene3D domains from tables produced by uniparc. |
| uniparc-domain-wstructure<br>(https://gitlab.com/datapkg/uniparc-domain-wstructure) | Scripts detailing the process of mapping of structural information in the PDB to Gene3D domain sequences extracted from UniParc. |

**Table S2.** Time performance comparison generating N designs using either ProteinSolver or Rosetta protocols (detailed in Table S3) for structures of different sizes. Average time in seconds for 7 independent runs on a single core of an Intel i7-5820K CPU @ 3.30GHz and an NVIDIA Titan V GPU. Enclosed in square brackets is the lowest REU produced. Lower boundary time estimation for Rosetta Optimized for N=1,000 was calculated by linear projection of N=1.

| PDB | N | # Residues | ProteinSolver | Rosetta optimized |
| --- | --- | --- | --- | --- |
| 5VLI | 1 | 39 | 0.168 s<br>[-90.87 REU] | 577 s<br>[-57.94 REU] |
| 1N5U | 1 | 92 | 0.400 s<br>[-234.53 REU] | 1,385 s<br>[-377.26 REU] |
| 4Z8J | 1 | 96 | 1.04 s<br>[-275.17 REU] | 1,742 s<br>[-161.325 REU] |
| 4UNU | 1 | 109 | 0.555 s<br>[-269.09 REU] | 1,963 s<br>[-174.46 REU] |
| 4BEU | 1 | 217 | 2.52 s<br>[-580.37 REU] | 4,547 s<br>[-375.83 REU] |

| PDB | N | # Residues | ProteinSolver | Rosetta default | Rosetta optimized |
| --- | --- | --- | --- | --- | --- |
| 5VLI | 1,000 | 39 | 96 s<br>[-119.15 REU] | 10,651 s<br>[-54.75 REU] | >280,000 s |
| 1N5U | 1,000 | 92 | 228 s<br>[-250.18 REU] | 28,843 s<br>[-133.22 REU] | >690,000 s |
| 4Z8J | 1,000 | 96 | 239 s<br>[-291.86 REU] | 34,068 s<br>[-152.78 REU] | >870,000 s |
| 4UNU | 1,000 | 109 | 271 s<br>[-298.21 REU] | 39,424 s<br>[-151.95 REU] | >980,000 s |
| 4BEU | 1,000 | 217 | 828 s<br>[-611.46 REU] | 100,795 s<br>[-346.98 REU] | >2,200,000 s |

**Table S3.**
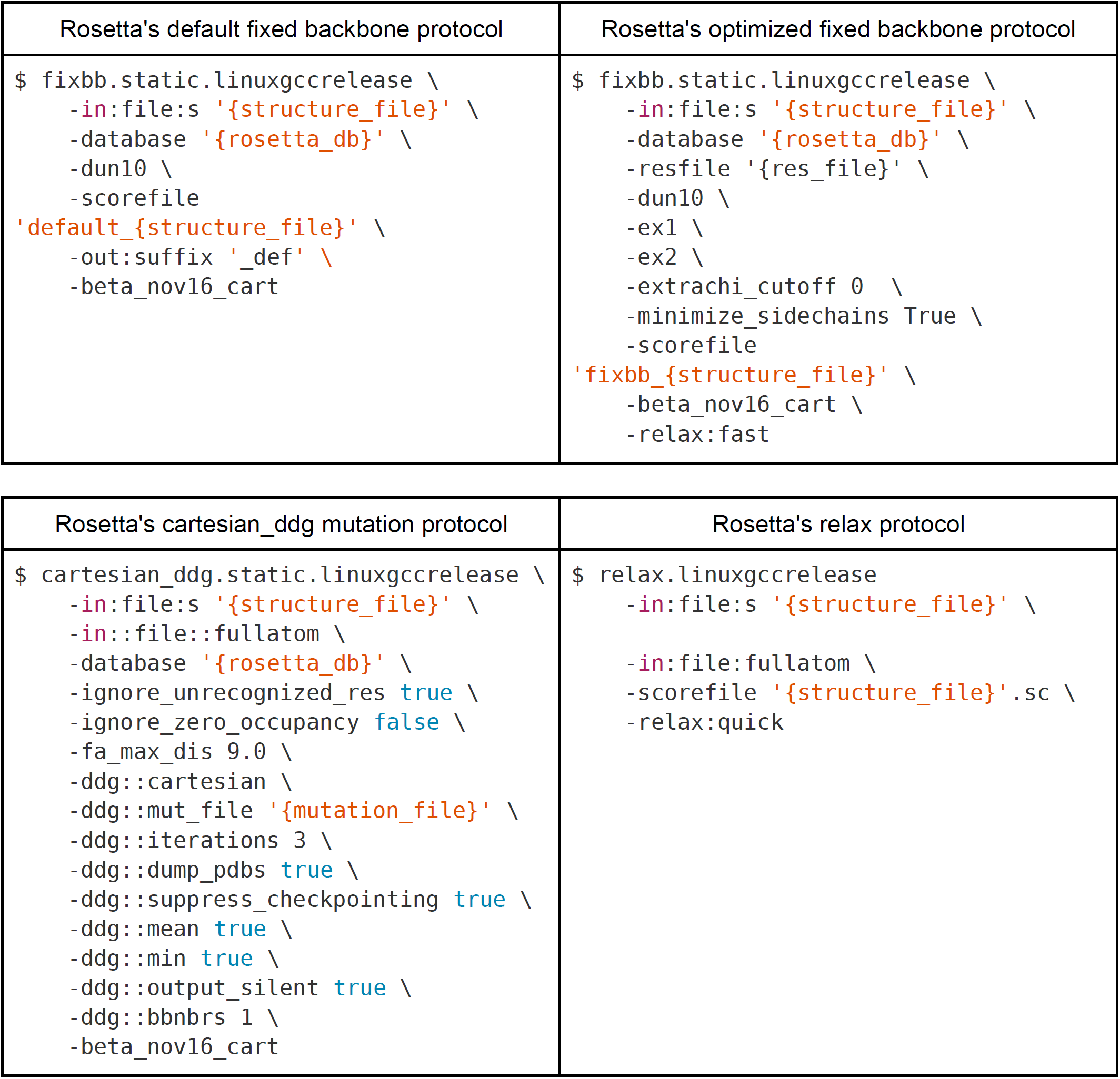
System commands used to run Rosetta’s default fixed backbone design protocol (upper left), Rosetta’s optimized fixed backbone design protocol (upper right), Rosetta’s “cartesian_ddg” mutation protocol (bottom left), and Rosetta’s relax protocol (bottom right).

**Table S4.** Similarity between sequences that have undergone successful experimental validation and sequences of the reference structures (**sequence identity to reference structure**) and the training database (**E-value for closest match in the training database; CATH superfamily of closest match in the training database**).

| **CATH domain** | **CATH superfamily** | **Generated sequence id** | **Sequence identity to reference structure** | **E-value for closest match in the training database** | **CATH superfamily of closest match in the training database** |
| --- | --- | --- | --- | --- | --- |
| 1n5uA03 | 1.10.246.10 | 1n5uA03-00 | 42.4% | 385.5 | 3.40.930.10 |
| 4beuA02 | 3.20.20.10 | 4beuA02-00 | 37.7% | 7.177 | 1.20.120.530 |

**Table S5.**
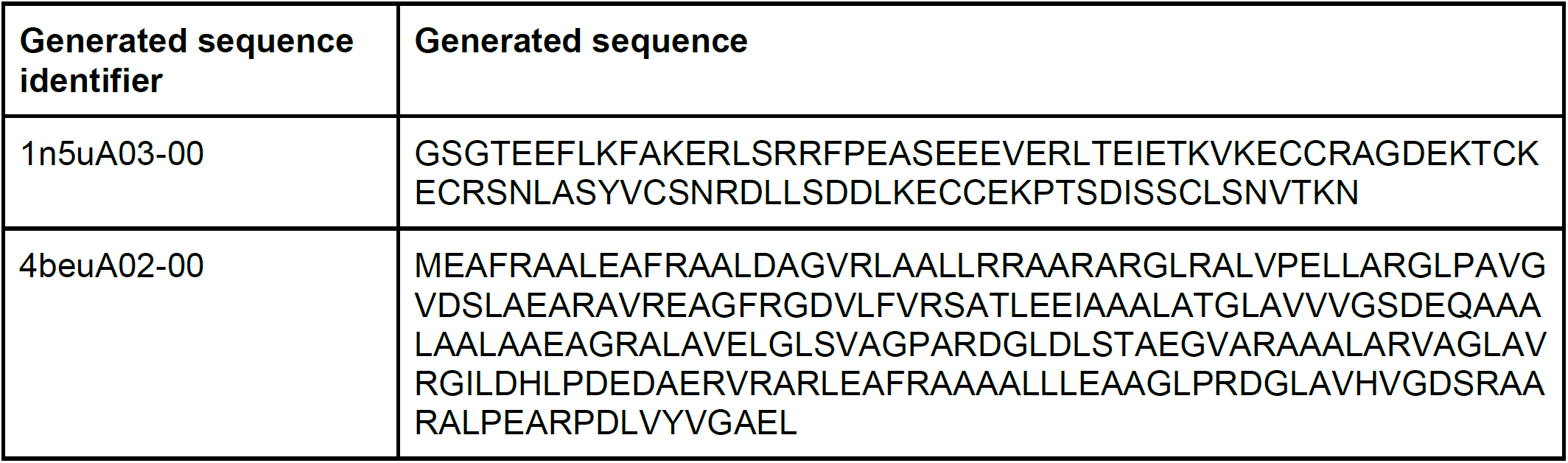
Sequences that have undergone successful experimental validation.

### Supplemental Figures

**Fig. S1.**
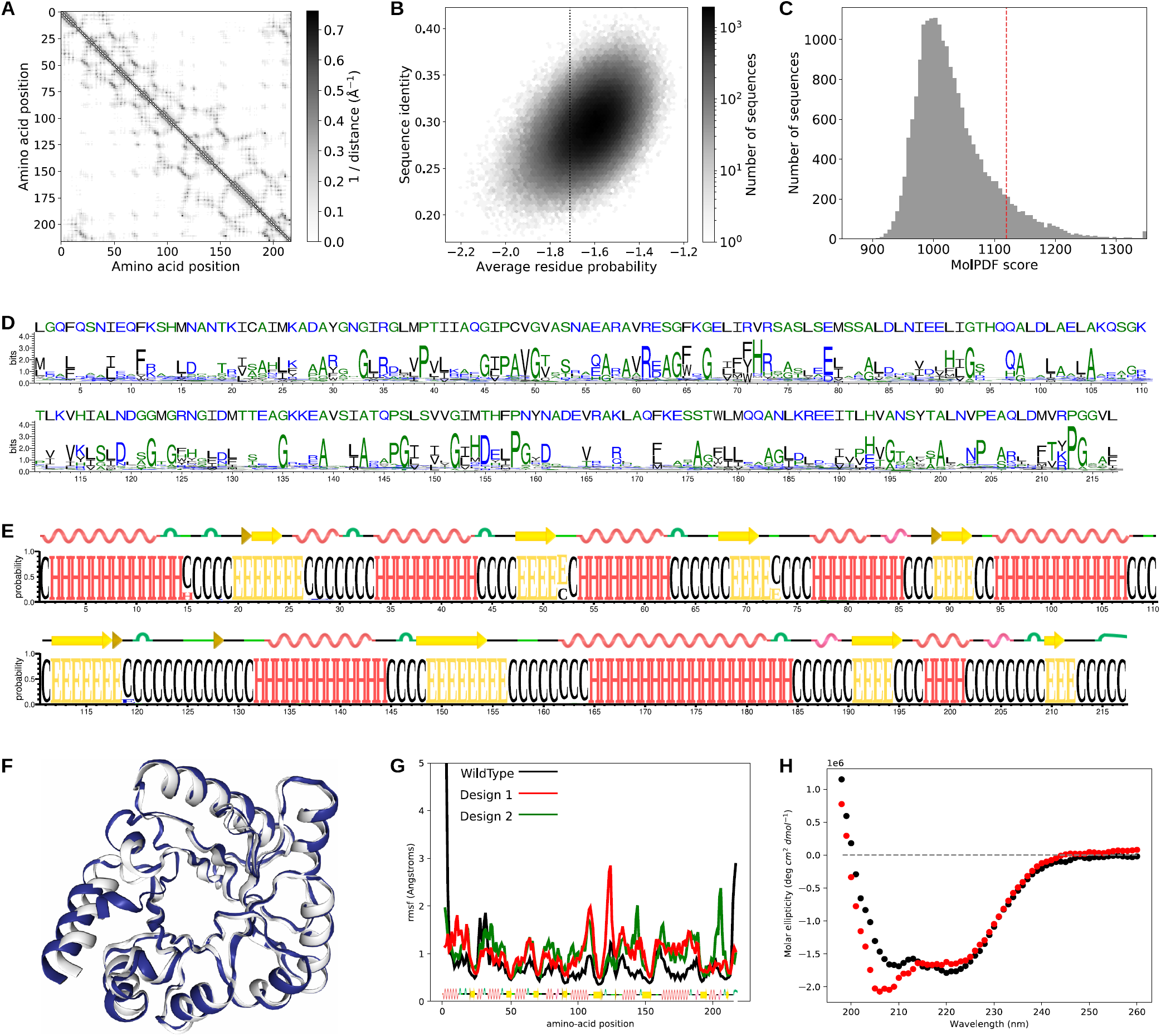
Computational and experimental validation of sequences generated to match the architecture of *alanine racemase*. **(A)** Amino acid contact map extracted from PDB structure *4BEU* and provided as input to the network. **(B)** Correlation between residue probabilities and sequence identities to the reference structure, for ∼600,000 sequences generated by the trained ProteinSolver network. **(C)** Distribution of Modeller MolPDF scores for homology models constructed for a sample of 20,000 generated sequences. **(D)** Sequence LOGO showing the conservation of residues in the generated sequences. **(E)** Secondary structure LOGO showing predicted secondary structures for a sample of 20,000 generated sequences. **(F)** Structure of the reference protein (white) overlaid with the structure of a model produced for one of the generated sequences using QUARK, a de-novo structural modelling tool (blue). **(G)** Average residue fluctuation in 100 ns molecular dynamics simulations of the reference structure and two homology models of generated sequences. **(H)** Circular dichroism spectra of the reference protein (black) and a generated protein (red).

**Fig. S2.**
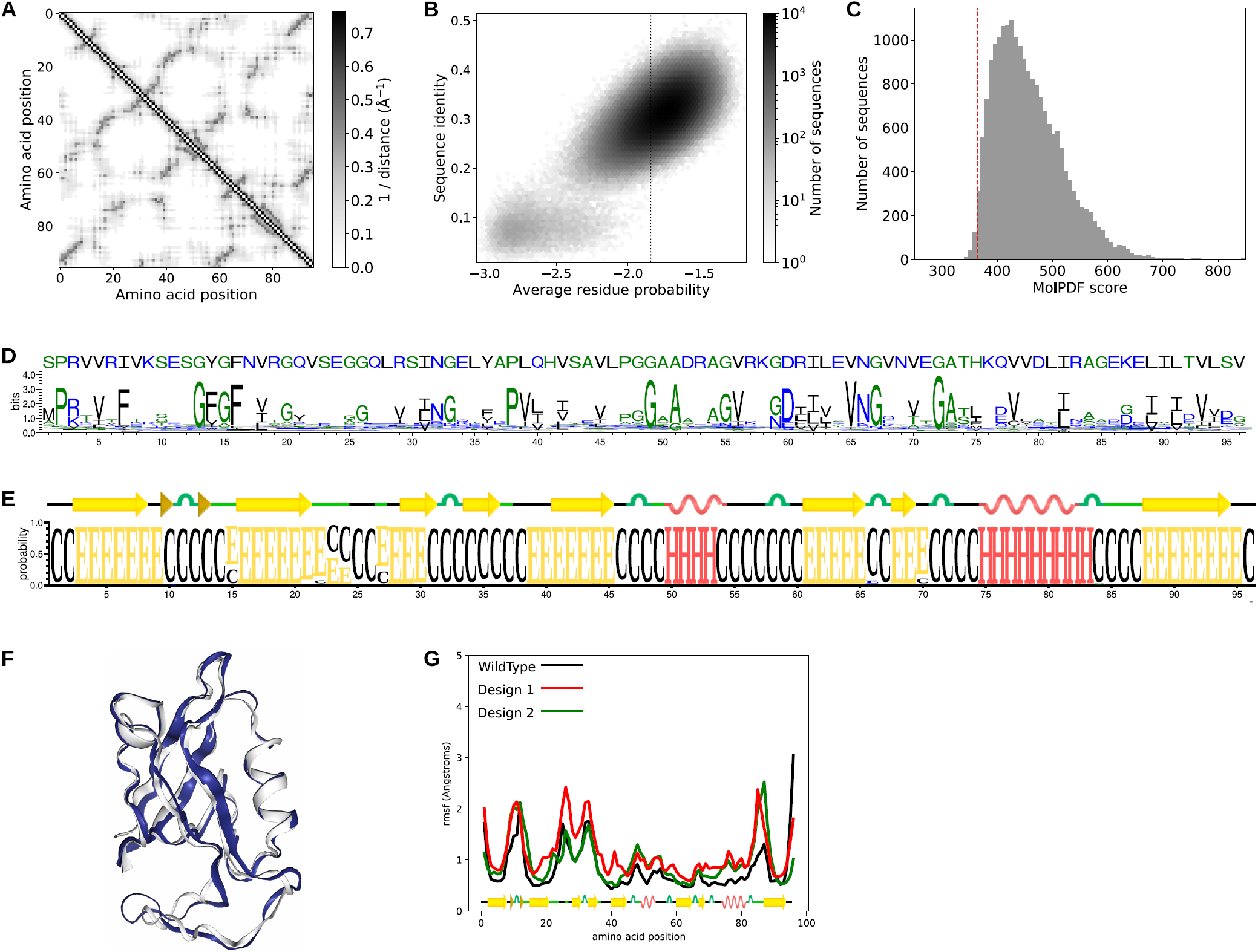
Computational and experimental validation of sequences generated to match the architecture of a *PDZ3* domain. **(A)** Amino acid contact map extracted from PDB structure *4Z8J* and provided as input to the network. **(B)** Correlation between residue probabilities and sequence identities to the reference structure, for ∼1.75 million sequences generated by the trained ProteinSolver network. **(C)** Distribution of Modeller MolPDF scores for homology models constructed for a sample of 20,000 generated sequences. **(D)** Sequence LOGO showing the conservation of residues in the generated sequences. **(E)** Secondary structure LOGO showing predicted secondary structures for a sample of 20,000 generated sequences. **(F)** Structure of the reference protein (white) overlaid with the structure of a model produced for one of the generated sequences using QUARK, a de-novo structural modelling tool (blue). **(G)** Average residue fluctuation in 100 ns molecular dynamics simulations of the reference structure and two homology models of generated sequences.

**Fig. S3.**
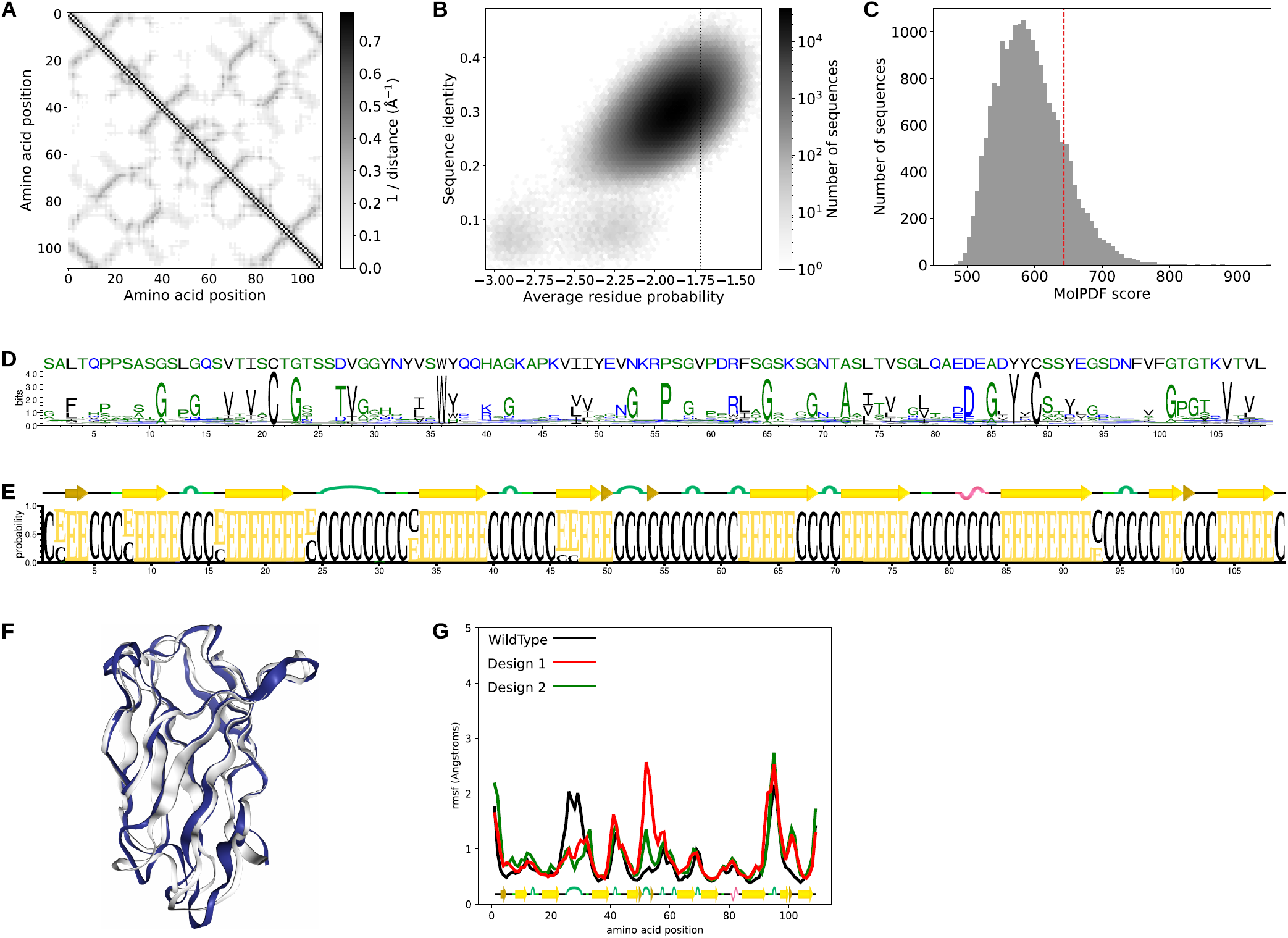
Computational and experimental validation of sequences generated to match the architecture of *immunoglobulin*. **(A)** Amino acid contact map extracted from PDB structure *4UNU* and provided as input to the network. **(B)** Correlation between residue probabilities and sequence identities to the reference structure, for ∼6.5 million sequences generated by the trained ProteinSolver network. **(C)** Distribution of Modeller MolPDF scores for homology models constructed for a sample of 20,000 generated sequences. **(D)** Sequence LOGO showing the conservation of residues in the generated sequences. **(E)** Secondary structure LOGO showing predicted secondary structures for a sample of 20,000 generated sequences. **(F)** Structure of the reference protein (white) overlaid with the structure of a model produced for one of the generated sequences using QUARK, a de-novo structural modelling tool (blue). **(G)** Average residue fluctuation in 100 ns molecular dynamics simulations of the reference structure and two homology models of generated sequences.

**Fig. S4.**
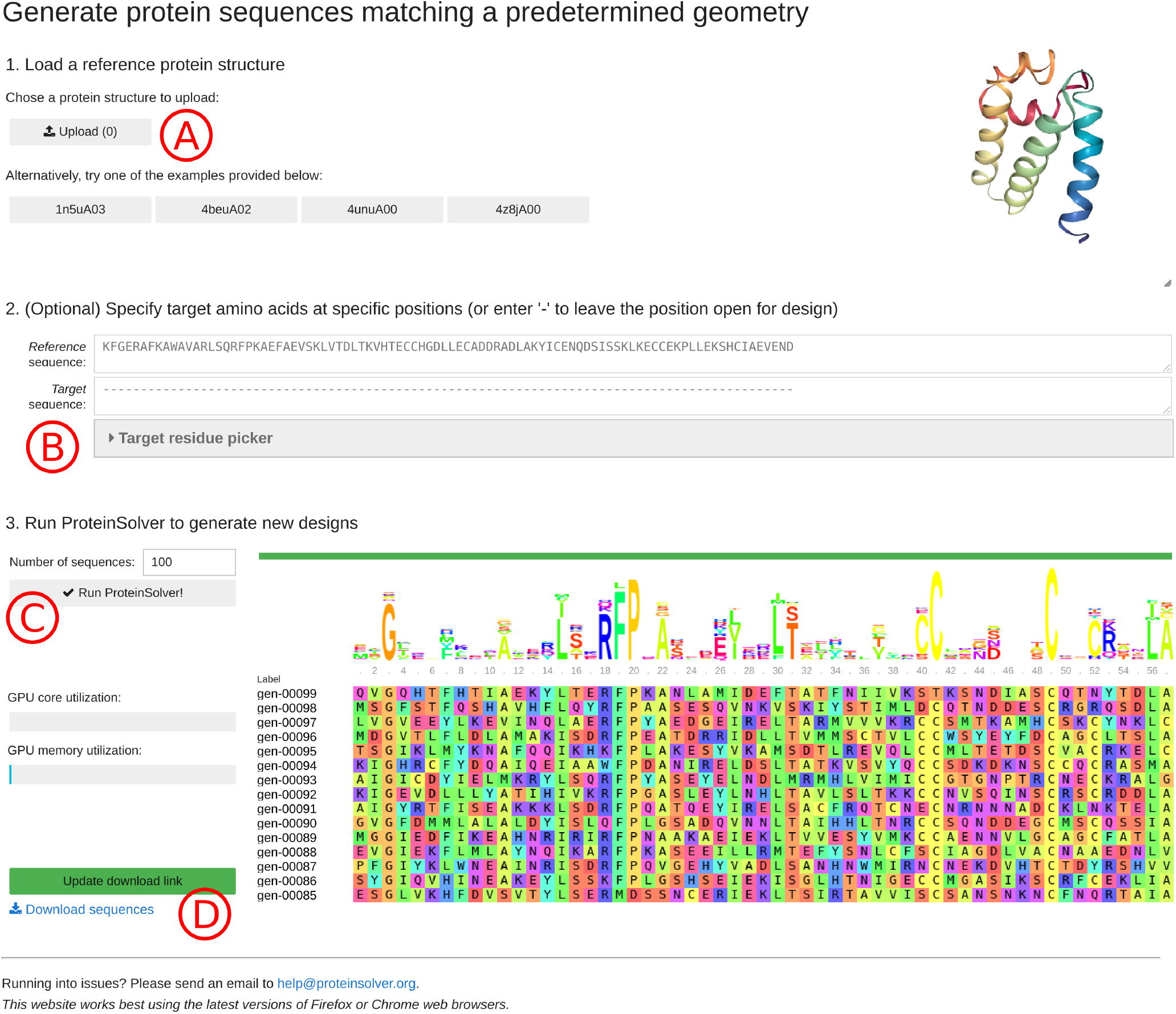
Screenshot of the ProteinSolver design web server (http://design.proteinsolver.org). **(A)** The user can upload the structure of a reference protein whose geometry will be used to restrain the space of generated amino acid sequences. Alternatively, the user can select one of four example proteins. **(B)** The user is given the option to explicitly fix one or more amino acids in the generated sequences to specific residues. **(C)** When the user clicks the “Run ProteinSolver” button, a background ProteinSolver process starts generating sequences matching the specified geometric and amino acid constraints. By default, 100 sequences are generated, although this number can be adjusted. The progress of the ProteinSolver process can be monitored by looking at the progress bar and the sequence logo displayed to the right of the “Run ProteinSolver” button, while GPU utilization can be monitored by looking at the status bars displayed below the “Run ProteinSolver” button. **(D)** Once a sufficient number of sequences have been generated, the user can click the “Generate download link” button, at which point a download link will appear, allowing the user to download the generated designs.

**Fig. S5.**
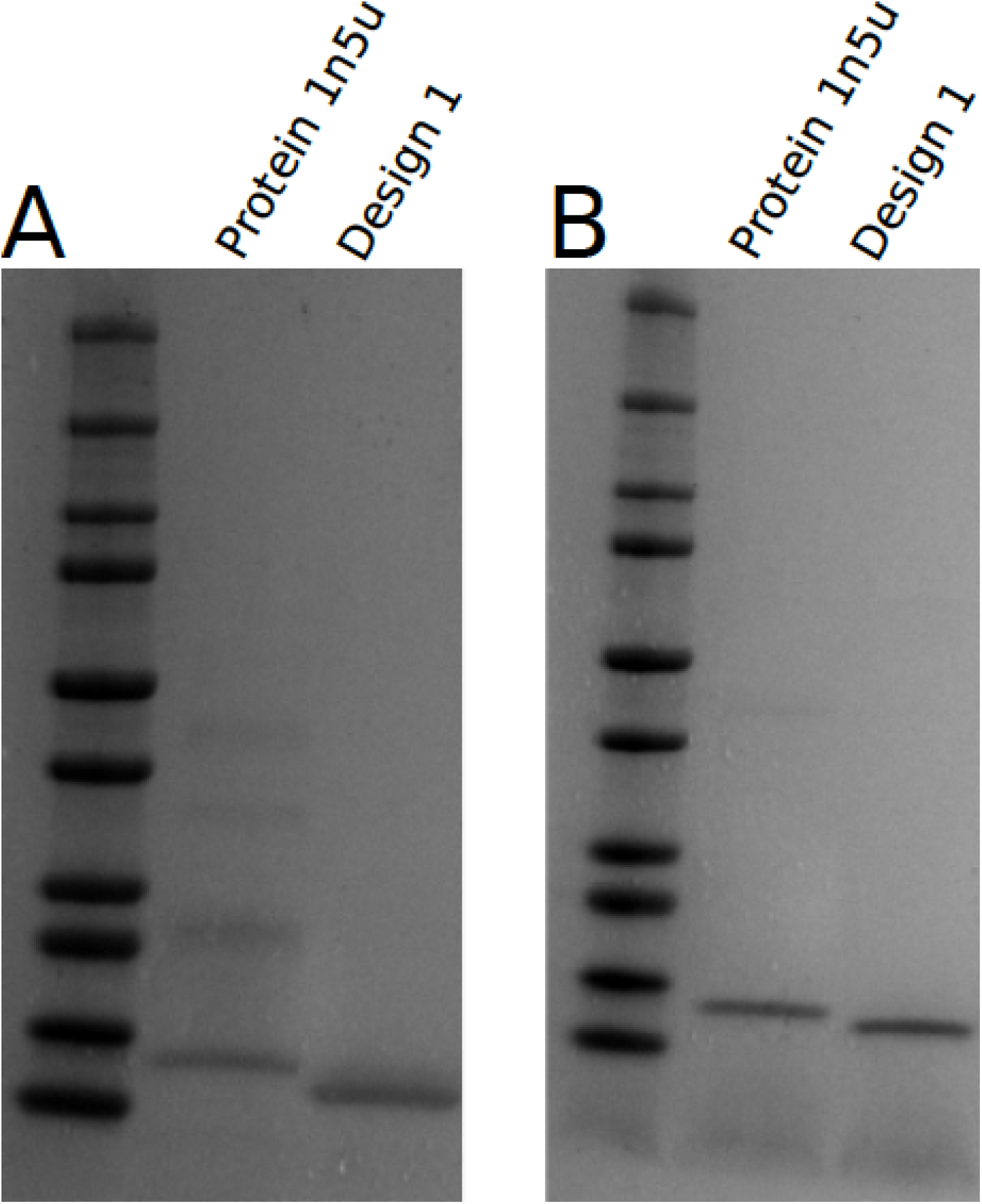
The designed albumin sequence and its natural counterpart (pdb-id: 1n5u) were analyzed by SDS-PAGE in **(A)** oxidizing and **(B)** reducing conditions (1 mM DTT). The natural protein, with an unpaired cysteine, forms high MW species not present in reducing conditions. Design 1 is only represented as a single band in both environments. Interestingly, the chosen sequence from the albumin template contains 4 pairs of potential disulfide bonds, whereas it is known that the albumin template used has only 3 of these bonds (PDB 1n5u). The designed 1n5u run in an SDS-PAGE in oxidising conditions as a single band. Furthermore, the mass spectrometry analysis by electrospray (ESI) of the molecular weight (MW) of designed 1n5u showed a loss of 8 Da against the theoretical MW, consistent with the loss of 8 protons. All together this indicates that a new disulfide bond not present in the albumin structural template was efficiently inserted into the designed sequence.

